# Vimentin intermediate filaments organize organellar architecture in response to ER stress

**DOI:** 10.1101/2022.03.24.485587

**Authors:** Tom Cremer, Lenard M. Voortman, Erik Bos, Daphne M. van Elsland, Laurens R. ter Haar, Roman I. Koning, Ilana Berlin, Jacques Neefjes

**Affiliations:** Department of Cell and Chemical Biology; Oncode Institute, Leiden University Medical Center LUMC, Leiden NL

**Keywords:** RNF26, Vimentin, intermediate filaments, endosomes, ER stress

## Abstract

Compartmentalization of organelles in space and time affects their functional state and enables higher order regulation of essential cellular processes. How organellar residence is maintained in a defined area of the cell remains poorly understood. In this study, we uncover a new role for intermediate filaments in the maintenance of organellar architecture and dynamics, which is executed through a functional connection between Vimentin and the ER-embedded ubiquitin ligase ring finger protein 26 (RNF26). While the ubiquitin ligase function of RNF26 promotes perinuclear positioning of endolysosomes, its catalytically inactive mutant I382R preferentially binds Vimentin through the RNF26 C-terminal tail. Loss of either RNF26 or Vimentin redistributes endolysosomes throughout the cytosol and mobilizes ER membranes from the perinuclear ER towards the periphery. Furthermore, RNF26 and Vimentin control changes in ER morphology and organelle compartmentalization during ER stress. Collectively, we define a new function for Vimentin-containing intermediate filaments as anchors of a dynamic interplay between the ER and endosomes, critical to the integrity of the perinuclear ER and corresponding perinuclear endosomal cloud during homeostatic and stress conditions.

**Synopsis:** The perinuclear area hosts a wide variety of cellular organelles, and their interaction with the ER governs essential cellular processes. To spatiotemporally organize endosomes and ER in the perinuclear region, the ER-embedded E3 ubiquitin ligase RNF26 interacts with Vimentin to physically link the perinuclear ER membrane with the intermediate filament cytoskeleton. As a result, Vimentin ensures perinuclear RNF26 retention, which in turn controls the perinuclear location of ER membranes and endosomes, which can be affected during stressed conditions.

- Vimentin interacts with inactive RNF26 in the ER membrane
- RNF26 by virtue of the Vimentin interaction controls perinuclear organization of ER membranes and the endosomal system
- Vimentin immobilizes ER membranes in the perinuclear area
- Vimentin and RNF26 compartmentalize organelles in the perinuclear region during ER stress
- We define a new function of Vimentin intermediate filaments in the control of the perinuclear endosomal and ER organization

## Introduction

In mammalian cells, the perinuclear region stages a get-together for diverse membranous organelles that collectively execute essential aspects of cellular biology (Cohen, Valm et al., 2018, Neefjes, Jongsma et al., 2017). Perinuclear localization of the ‘rough’ endoplasmic reticulum (ER), Golgi, mitochondria, and endolysosomes support cellular processes such as energy production, control of nutrient sensing and proteostasis (Johnson, Ostrowski et al., 2016, Korolchuk, Saiki et al., 2011, Leitman, Ron et al., 2013, Starling, Yip et al., 2016), modulation of signaling responses (Cremer, Jongsma et al., 2020a, Jia & Bonifacino, 2019) and maintenance of cell polarity (Ang & Fölsch, 2012). It is well appreciated that long-range transport of organelles to the perinuclear region is typically driven by the microtubule-associated dynein motor (Cabukusta & Neefjes, 2018). However, what maintains organelles at their site of function is often less clear. Accumulating evidence indicates that, in addition to cytoskeletal determinants and motor-based transport, intracellular distribution of organelles is guided by specialized membrane contact sites (MCSs) formed between autonomous compartments (Phillips & Voeltz, 2016, Prinz, Toulmay et al., 2020). While the sheer crowdedness of the perinuclear cytoplasm makes a wide range of cross-compartmental interactions possible, how these different elements come together to finetune organellar localization, dynamics, and ultimately function remains largely unknown. In the present study, we identify a key role for intermediate filaments (IFs) as perinuclear anchors for the ER and the endolysosomal system and reveal a dynamic molecular interplay driving co-organization of these compartments in response to ER stress.

The ER is the cell’s most expansive organelle (Goyal & Blackstone, 2013) that masterfully coordinates cellular homeostasis. To facilitate its breadth of function, the ER network is divided into biochemically and morphologically distinct regions (Leitman et al., 2013, Nixon-Abell, Obara et al., 2016). The densely packed sheet-like perinuclear ER segment functions in protein biosynthesis, folding and quality control, and forms the front line of stress response, while the dynamic peripheral ER network manages calcium sensing and uptake (Cremer, Neefjes et al., 2020b). Furthermore, to influence processes occurring elsewhere in the cell, the ER engages virtually all other organelles in a diverse array of MCSs (Wu, Carvalho et al., 2018). Juxtaposition of membranes at these contacts, driven by reversible *in trans* pairings of dedicated tethers, provides unique opportunities for interorganellar communication and material exchange (King, Sengupta et al., 2020, Prinz et al., 2020). Specifically, MCSs between the ER and the endolysosomal system have been extensively documented to play important roles in lipid homeostasis (Jain & Holthuis, 2017) and calcium signaling (Cremer et al., 2020b). ER MCSs are also directly involved in decisions pertaining to transport (Raiborg, Wenzel et al., 2015, Rocha, Kuijl et al., 2009, Saric, Freeman et al., 2021), fusion (Wijdeven, Janssen et al., 2016), and fission (Hoyer, Chitwood et al., 2018) of endosomes and lysosomes. In return, endosomes facilitate ER movement along microtubule tracks and thereby support continuous remodeling of the ER network (Lu, van Tartwijk et al., 2020, Spits, Heesterbeek et al., 2021). This ER hitchhiking phenomenon occurs largely on late endosomes (Spits et al., 2021), which engage the ER with greater frequency and residence time relative to their early or less mature counterparts (Friedman, Dibenedetto et al., 2013). Under steady state conditions, the bulk of endolysosomes congregates in the perinuclear cytoplasm, and the architecture and dynamics of this perinuclear vesicle cloud are regulated by the ER-embedded ubiquitylation complex comprised of the E3 ligase RING finger protein 26 (RNF26) and its cognate conjugating enzyme UBE2J1 (Cremer et al., 2020a, Jongsma, Berlin et al., 2016). RNF26 resides in the perinuclear ER membrane (Fenech, Lari et al., 2020, Jongsma et al., 2016) and this feature enables it to spatially restrict various endolysosomal species by attracting their associated adaptor proteins in a manner dependent on its ubiquitylation activity (Jongsma et al., 2016). However, what ensures perinuclear retention of RNF26 and other proteins within the greater ER membrane is unclear. Moreover, whether RNF26 and its molecular partners contribute to the global governance of ER architecture and dynamics has not been explored.

In this study, we identify Vimentin—a major constituent of IFs—as a perinuclear anchor for RNF26 and hereby uncover a new molecular bridge between the ER and the IF cytoskeleton. We show that both RNF26 and Vimentin promote maintenance of the perinuclear ER and coordinate changes in its morphology. We observe that Vimentin preferentially binds inactive RNF26, restricting it within the perinuclear ER segment. Subsequent release of RNF26 upon activation then enables corresponding spatial arrangement of endolysosomes. Through this dynamic interplay, Vimentin and RNF26 guard organellar distribution in steady state and drive acute perinuclear coalescence of the ER with endolysosomes in response to ER stress. Type III IFs composed of Vimentin have thus far been appreciated predominantly for their structural contributions to cell rigidity (Etienne-Manneville, 2018, Lowery, Kuczmarski et al., 2015, Toivola, Tao et al., 2005) and as markers of the epithelial-to-mesenchymal transition during cancer progression (Ivaska, 2011, Usman, Waseem et al., 2021). Our findings now position IFs, along with the microtubule and actin networks (Svitkina, 2018, Zheng, Obara et al., 2022), as dynamic regulators of cellular compartmentalization under steady state and mobilization of organelles in response to stress.

## Results

### RNF26 associates with the Vimentin cytoskeleton

Under steady state conditions, adherent cells display distinct organellar distribution between their rather sparse and highly dynamic periphery (PP) and a crowded perinuclear (PN) region (Fig. EV1A, PP vs PN). This PN/PP dichotomy is reflected by the organization of the ER network, which features interconnected peripheral tubules and densely packed membranes residing near the nucleus (Fig. 1A). Proteolytic compartments belonging to the endolysosomal system also congregate in the perinuclear space (Fig. 1A), and this arrangement depends on the RING activity of the ER-embedded ubiquitin ligase RNF26 (Jongsma et al., 2016). These considerations led us to hypothesize that compartmentalization of the ER and the endolysosomal system may be coregulated through RNF26. We have previously found that full length RNF26 concentrates in the PN ER segment, congruent with its facilitation of the PN vesicle cloud (Jongsma et al., 2016). By contrast, RING deletion mutant of RNF26 distributes throughout the ER, implying active PN retention of otherwise freely diffusing RNF26 within the ER membrane. We therefore considered whether interactors of the RING domain of RNF26 include factors driving its retention within the PN ER membrane. These same factors could then also be expected to define the identity of the PN ER subdomain and influence intracellular distribution of endosomes. Among proteins co-precipitating with the RING domain of RNF26 we identified Vimentin (Jongsma et al., 2016), which assembles into PN intermediate filaments frequently overlapping with ER membranes (Fig. 1A, B). Remarkable similarity in distribution between Vimentin filaments and the PN ER by immunofluorescence prompted us to observe them at higher resolution (Fig. EV1B). 3D reconstructions of TEM tomograms collected from a PN cell region revealed connections between ER membranes and IFs, either directly or spaced by protein condensates (Fig. 1C, zoom-ins, Fig. EV1C, Movie 1). This indicated that the Vimentin cytoskeleton could be a viable candidate for anchoring the perinuclear ER, and by extension its interacting organelles.

**Figure 1:**
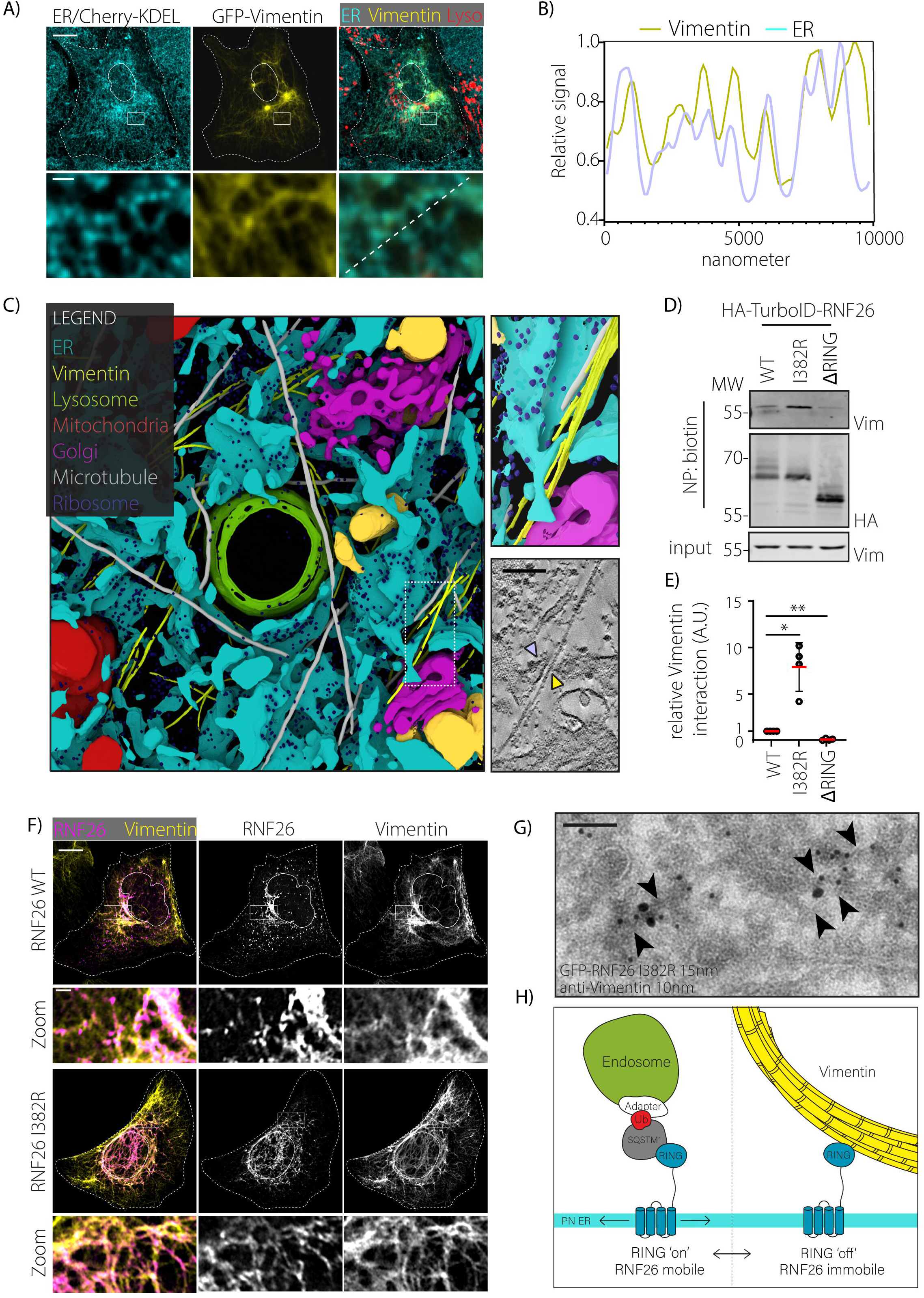
ER-embedded RNF26 interacts with Vimentin in the perinuclear area. **A)**Intracellular distribution of the ER and Vimentin cytoskeleton. A representative confocal image of U2OS cells transiently transfected to express ER-targeted RFP-KDEL (cyan) GFP-Vimentin (yellow) is shown. Merged image also includes SiR-Lysosome staining (red) designating proteolytic endolysosomes. Zooms highlight coalescence of ER membranes with Vimentin cytoskeleton in the perinuclear (PN) region. Cell and nuclear boundaries are demarcated using dashed and continuous lines, respectively. Scale bar = 10μm, zoom scale bars = 1 μm. **B)** Line intensity plots of GFP (ER) and RFP (Vimentin) signal along the dotted line in (A). **C)** 3D rendering of a PN area from a U2OS cell, constructed from tomograms of three serial EM sections from the area designated in *Fig. EV1B*. Manual color annotation of organelles is provided in the legend. Boxed zoom-ins of renders and tomogram slices highlight a long bundle of filaments juxtaposed to ER membranes. *See also Fig. EV1B*. Scale bar = 100 μm **D)** Contribution of ubiquitin ligase RING domain determinants to the *in situ* interaction between RNF26 and Vimentin. Proximity biotinylation by 2HA-TurboID-RNF26 wild-type (WT), catalytically inactive point mutant (I382R), or RING domain truncation mutant (ΔRING) in HeLa cells following 30 min incubation with 100μM biotin prior to lysis and neutravidin precipitation (NP). Immunoblot analysis of precipitates and their corresponding whole cell lysates (WCL) against HA and endogenous Vimentin is shown. Molecular weight markers indicated. **E)** Quantifications of Vimentin biotinylation by 2HA-TurboID-RNF26 and variants from (B) normalized to autobiotinylation of HA-tagged species and expressed relative to RNF26-WT; n= 4 independent biological replicates per condition, error bars correspond to +/− SD (t-test. ** p<0,01; *** p<0,001; **** p<0.0001). **F)** Localization of RNF26-positive ER membranes along Vimentin filaments as a function of ubiquitin ligase activity. Near-superresolution confocal fluorescence images of U2OS cells ectopically expressing RFP-RNF26 (WT versus I382R), fixed and immunostained against endogenous Vimentin. Single channel images and color overlays are shown. Boxed zoom-ins highlight PN regions. Cell and nuclear boundaries are demarcated using dashed and continuous lines, respectively. Scale bar = 10μm, zoom scale bars = 1 μm. **G)** Juxtaposition of ER membranes to Vimentin filaments. Immuno-EM micrograph of U2OS cell transiently overexpressing GFP-RNF26 I382R and labeled for GFP (15nm gold) and endogenous Vimentin (10nm gold). Arrows indicate ER structure. **H)** Model on RNF26/Vimentin interaction, distilled from Fig. 1, Fig. S1. and (Jongsma et al., 2016). RNF26 may appear in two distinct enzymatic states, either enzymatically active and prone to form the endosomal positioning complex in conjunction with SQSTM1 and vesicle adapters as previously described (Jongsma *et al*., 2016), or enzymatically inactive to interact with the Vimentin cytoskeleton. While active RNF26 may move freely in the ER lipid bilayer, inactivity immobilizes it in the ER membrane through interaction with Vimentin. Cell and nuclear boundaries are demarcated using dashed and continuous lines, respectively.

Using proximity labeling (Branon, Bosch et al., 2018), we verified the interaction between Vimentin and full-length wild type (WT) RNF26 fused to a promiscuous biotin ligase moiety TurboID (Fig. 1D, E). As expected, the complex with Vimentin was no longer detected upon removal of the RING domain from RNF26 (Fig. 1D, E). Since RING domains impart enzymatic activity to E3 ubiquitin ligases, we also assessed the catalytically inactive RNF26-IR point mutant for its interaction with Vimentin. Surprisingly, we found that Vimentin interaction was markedly enhanced by inactivation of RNF26 ubiquitin ligase function (Fig. 1D, E). We have previously shown that ubiquitylation by RNF26 is mediated in conjunction with the E2 ubiquitin conjugating enzyme UBE2J1 (Cremer et al., 2020a). Its ER-based homologue UBE2J2 is a pseudo-compatible E2 for RNF26, and its overexpression renders the enzymatic complex inactive (Cremer et al., 2020a). As a result, the RNF26/UBE2J2 complex exhibited improved binding to Vimentin (Fig. EV1D), suggesting that regulation of RNF26 ligase activity by E2 enzyme availability affects connections between the ER and the IF cytoskeleton.

We further noted that RNF26-WT contacts the Vimentin IF cytoskeleton in a punctate fashion, while the IR mutant displays a filamentous appearance and aligns with Vimentin filaments in the PN region of the cell (Fig. 1F). Importantly, immuno-EM analysis demonstrated that RNF26-IR remains in the ER membrane when in proximity to Vimentin (Fig. 1G). These results suggest that inactivity of RNF26 instigates persistent association with Vimentin filaments. In line with this, FRAP experiments conducted to measure the behavior of RFP-RNF26 in real time point to reduced membrane mobility of inactive RNF26 as compared to either its catalytically competent counterpart or the RING deletion mutant (Fig. EV1E, F) that lacks the ability to bind Vimentin (Fig. 1D, E). Altogether, these results uncover a molecular link between the Vimentin cytoskeleton and RNF26 that may be controlled through regulation of RNF26 E3 ligase activity (Fig. 1H).

### Vimentin controls PN distribution of RNF26 and endolysosomes

Having identified Vimentin as an interactor of RNF26, we proceeded to investigate whether Vimentin influences retention of RNF26 in the PN ER membrane. To this end, we employed genome editing by CRISPR/Cas9 to create Vimentin knockout cells (Fig. 2A, Vim KO). Upon Vimentin ablation, RNF26-WT was found to redistribute towards the peripheral ER, as measured by overlap with the ER marker VAP-A (Fig. 2B, C). Meanwhile, RNF26-IR lost its perinuclear filamentous appearance and formed peripheral aggregate-like clusters, which were rescued by reintroduction of Vimentin into the knockout cells (Fig. EV2A-C). These results indicate that Vimentin is needed for normal distribution of RNF26 in both enzymatically active and inactive states.

**Figure 2:**
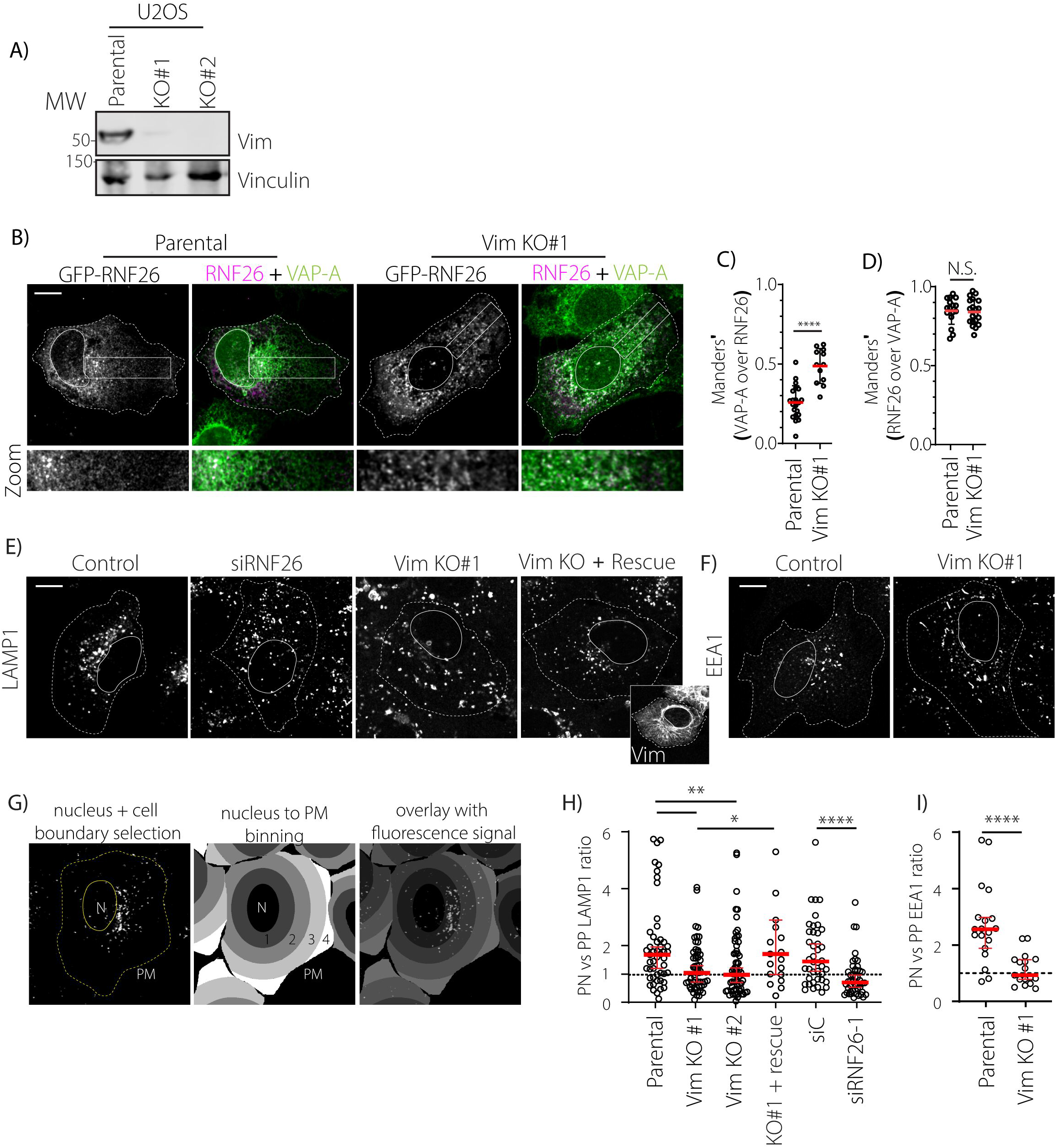
Vimentin is required for perinuclear localization of RNF26 and cognate endolysosomes. **A)** Validation of Vimentin knockout in U2OS cells. Western blot analysis of two Vimentin knockout U2OS cells created using CRISPR/Cas9. Membranes were stained for endogenous Vimentin, while Vinculin was stained as a loading control. MW markers indicated. **B)** Distribution of RNF26 as a function of Vimentin. Representative confocal microscopy images of overexpressed GFP-RNF26 distribution in WT and VIM KO#1 U2OS, immunostained for VAP-A to visualize the ER.. Boxed zoom-ins designate region spanning from the perinuclear area (PN) towards the plasma membrane (periphery, PP). Cell and nuclear boundaries are demarcated using dashed and continuous lines, respectively. Scale bar = 10μm. **C)** Colocalization (Manders’) of VAP-A over RNF26 in WT vs VIM KO#1 cells, indicating the fraction of the ER that is covered by RNF26. Error bars indicate mean +/− SD (t-test, **** p<0,001, *n* = 22 cells (WT) vs 12 cells (KO). **D)** Colocalization (Manders’) of RNF26 over VAP-A in WT vs VIM KO#1 cells, indicating presence of RNF26 in the ER after VIM KO. Error bars indicate mean +/− SD (t-test, N.S. = non-significant). **E, F)** Late (E) and early (F) endosomal distribution as a function of Vimentin (and RNF26 (E)). Shown are Z-slices of representative parental U2OS cell, siRNF26#1 (E), VIM KO#1 cells, and VIM KO#2 cells that were transfected with Vimentin. Cells were fixed and immunostained for LAMP1 (E) (and Vimentin (rescue, E) or EEA1 (F) prior to imaging with a confocal microscope. Cell and nuclear boundaries are demarcated using dashed and continuous lines, respectively. Scale bar = 10μm. **G)** Strategy for quantification of perinuclear vs peripheral fluorescent signals used in this paper. Briefly, cell nucleus and PM were manually annotated after which the cytoplasm was classified into 4 bins ranging from nucleus to PM. Then, using a custom Fiji plugin, fluorescent signals were merged with the binned image and mean fluorescence per bin and bin size were calculated. Average intensity per pixel in each bin was calculated, weighed for bin size and compared among bin 1 (PN) and bin 3 (PP) to obtain a ratio of perinuclear vs peripheral signal. **H)** Endosomal distribution analysis of late (LAMP1) and early (EEA1) endosomal distributions using Fiji plugin as described in (G). 51 (WT), 55 (VIM KO#1), 65 (VIM KO#2), 19 (VIM KO#1 + rescue), 42 (siC) and 48 (siRNF26) cells were analyzed (Mann-Whitney U test, (* p<0,05; ** p<0,01; **** p<0,0001). Error bars indicate median and 95% confidence interval

Since RNF26 regulates the architecture of the endolysosomal system, we next investigated the impact of Vimentin ablation in this context. Phenocopying RNF26-silenced cells, Vimentin knockout cells exhibited dispersed endolysosomes characterized by the LAMP1 marker, and this phenotype was also rescued by ectopic expression of Vimentin (Fig. 2E). Depletion of Vimentin similarly affected distribution of early endosomes marked by EEA1, indicating that, as in the case of RNF26, Vimentin also enforces spatial constraints across the endocytic repertoire (Fig. 2F). To quantify vesicular dispersion in an unbiased manner, we developed a Fiji plugin that calculates the distance between cytoplasmic fluorescent signals and the nucleus (Fig. 2G). Using this approach, concentration of vesicles in the endolysosomal cloud in unperturbed cells translates into a high signal ratio for bin 1 (PN) versus bin 3 (PP). Meanwhile, loss of either Vimentin or RNF26 results in a more homogeneous distribution of late and early endosomal signals throughout the cell, with a median PN/PP ratio approaching 1 (Fig. 2H). Taken together, these data indicate that Vimentin controls PN localization of RNF26 in the ER, and along with it, informs the PN accumulation of the endolysosomal system.

### C-terminal tail of RNF26 is necessary for Vimentin interaction

To investigate the molecular details of RNF26 retention within the PN ER, we aimed to pinpoint the region in RNF26 responsible for Vimentin binding. Structure prediction tools suggested that the segment C-terminal to the RING domain of RNF26 contains a β-sheet followed by a three-residue extension, which may potentially be compatible with protein-protein interactions. We therefore investigated whether Vimentin binding is affected by C-terminal truncation mutants (WT/Δ423-432 and IR/Δ423-432) of RNF26 (Fig. 3A). Strikingly, loss of the ultimate 10 C-terminal residues of RNF26 abrogated its interaction with Vimentin (Fig. 3B). In parallel, these mutants were found to mislocalize outside of the PN ER, displaying distributions similar to that of RNF26 in Vimentin KO cells (Fig. 2B). The same RNF26 variants also exhibited decreased colocalization with Vimentin filaments relative to full-length RNF26 (Fig. 3C, D) but maintained general residence in the ER membrane (Fig. 3E, Fig 2A, EV2A, EV3A). These results indicate that RNF26 requires its C-terminal tail to interact with Vimentin and localize to the correct ER subdomain. Examination of the composition of the RNF26 C-terminus revealed that the ultimate residues are evolutionary conserved among vertebrates and may thus be essential for this protein’s function(s) (Fig. 3F). Furthermore, the C-terminal tyrosine residue Y432 was previously shown to be essential for RNF26 activity (Fenech et al., 2020). Therefore, we wondered if this ultimate tyrosine residue is involved in binding RNF26 to Vimentin. Mutation of tyrosine 432 to alanine (Y432A) indeed abolished (auto)-ubiquitylation on RNF26 (Fig. EV3B) and partly abrogated Vimentin interaction as compared to the I382R inactive mutant (Fig. 3G). These results indicate that the ultimate tyrosine residue promotes both enzymatic activity and Vimentin binding capacity of RNF26.

**Figure 3:**
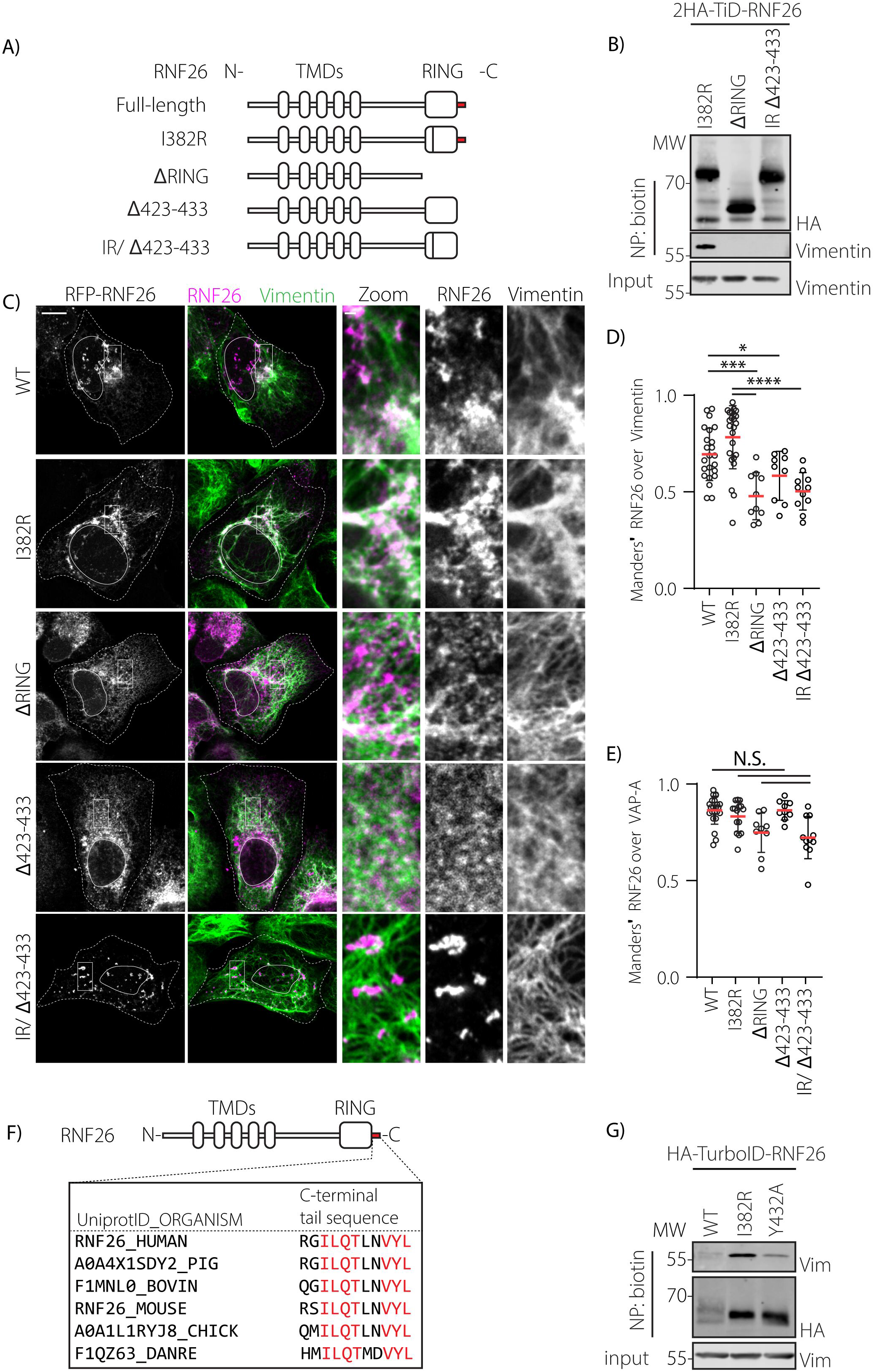
RNF26 interacts with Vimentin through its C-terminal tail. **A)** Schematic representation of RNF26 variants used in this study. Indicated are N-terminal transmembrane domains, the C-terminal RING domain and the I382R mutant herein, and C-terminal tail (in red). **B)** Interaction of RNF26 and Vimentin as a function of RNF26 truncation. Proximity biotinylation by 2HA-TurboID-RNF26 wild-type (WT) or RING domain truncation mutants (ΔRING and IR/Δ422-432) in HeLa cells following 30 min incubation with 100μM biotin prior to lysis and neutravidin precipitation (NP). Immunoblot analysis of precipitates and their corresponding whole cell lysates (WCL) against HA and endogenous Vimentin is shown. Molecular weight markers indicated. **C)** Intracellular distribution of RNF26 as a function of RNF26 activity and length described in (A). Representative Z-slice of U2OS cells that were transfected with RNF26 constructs, immunostained for Vimentin prior to imaging with a confocal microscope. Cell and nuclear boundaries are demarcated using dashed and continuous lines, respectively. Scale bars = 10μm, zoom scale bar = 1μm. **D)** Colocalization (Manders’) analysis of RNF26 over Vimentin from images in (C) (Students’ T-test. * p<0,05; ** p<0,01; *** p<0,001; **** p<0.0001). **E)** Colocalization (Manders’) analysis of RNF26 over VAP-A from images in Fig. 2B, EV2A and EV3A (Students’ T-test, N.S. non-significant). **F)** Primary structure homology analysis of ultimate C-terminal residues of RNF26 in human, pig, bovine, mouse, chicken and zebrafish species. Red residues are evolutionary conserved among all investigated species. **G)** Interaction of RNF26 and Vimentin as a function of RNF26 Y432 identity. Proximity biotinylation by 2HA-TurboID-RNF26 wild-type (WT) or inactive (I382R or Y432A) mutants HeLa cells following 30 min incubation with 100μM biotin prior to lysis and neutravidin precipitation (NP). Immunoblot analysis of precipitates and their corresponding whole cell lysates (WCL) against HA and endogenous Vimentin is shown. Molecular weight markers indicated.

### RNF26 and Vimentin regulate ER architecture in space and time

Like the endolysosomal system, the ER network is organized bilaterally, with densely packed perinuclear membranes and discrete more dynamic tubules in the cell periphery (Fig. 1, Fig. 4A). Meanwhile, Vimentin filaments also accumulate in the PN region, where they bind RNF26 (Fig. 1), prompting the question of whether this interaction influences the bilateral organization of the ER. Dramatically, depletion of either Vimentin or RNF26 was found to deteriorate the PN/PP dichotomy of the ER, hallmarked by dilution of VAP-A-positive membranes into the cell periphery (Fig. 4A, B, EV4A). This suggested homogenization of ER membranes in the absence of RNF26 or Vimentin. Since the PN ER is typically rich in ER sheets (Shibata, Voeltz et al., 2006, Terasaki, Shemesh et al., 2013), we also investigated the distribution of ER sheet marker CLIMP63 (Zheng et al., 2022). Once again, removal of RNF26 or Vimentin resulted in expansion of the CLIMP63-positive ER segment (Fig. 4C, EV4B). Collectively, these results suggest that RNF26 and Vimentin are required to maintain compartmentalization of membrane composition in the ER.

**Figure 4:**
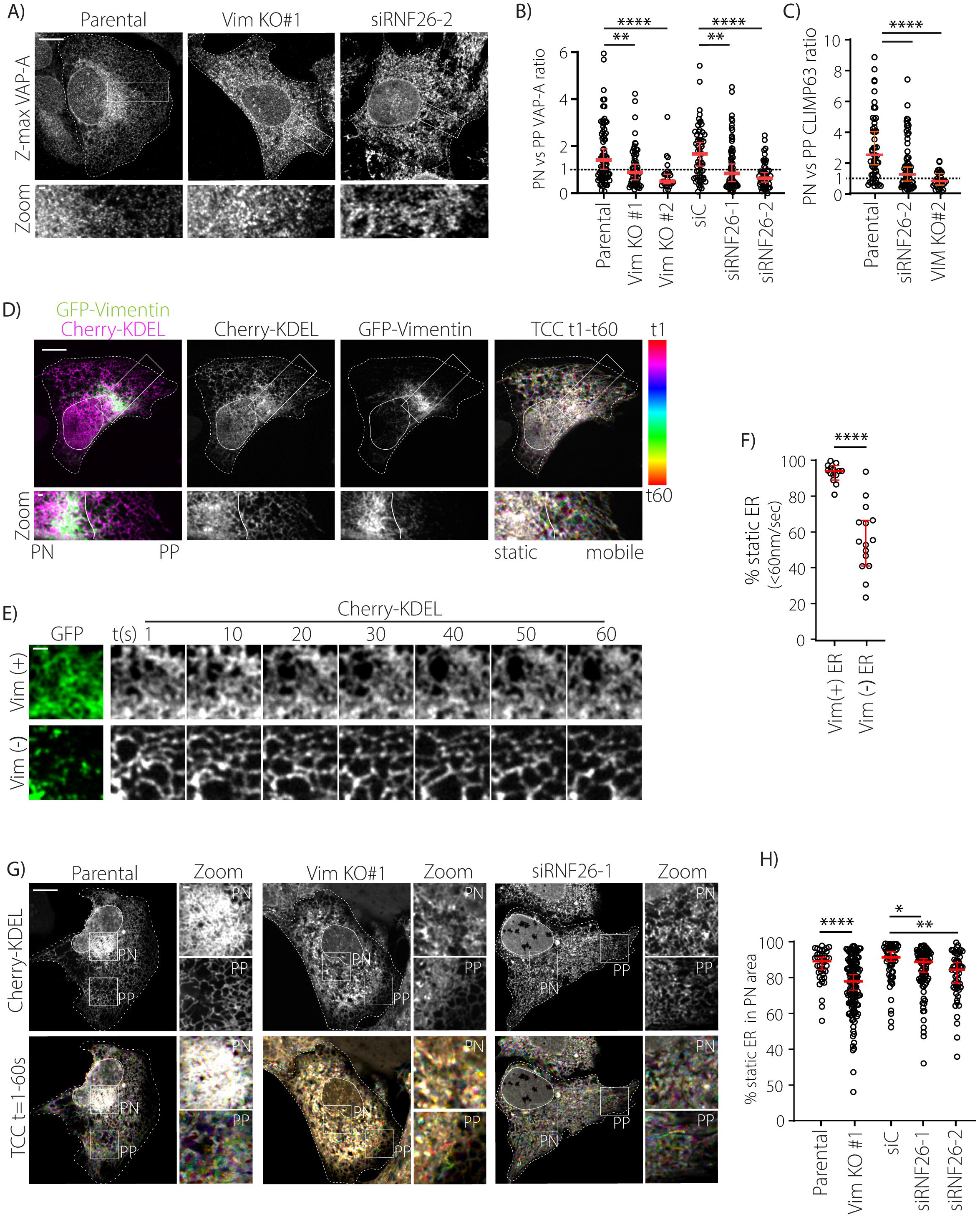
Interplay between Vimentin and RNF26 maintains integrity of the perinuclear ER. **A)** ER distribution as a function of RNF26 or Vimentin. Confocal analysis of parental U2OS cells, U2OS Vim KO #1 cells, or U2OS cells silenced for RNF26 (si#2) that were fixed and immunostained for VAP-A before Z-stack collection. Shown are maximum projections of representative cells. Zoom boxes show a select region spanning from the nucleus to the PM. Cell and nuclear boundaries are demarcated using dashed and continuous lines, respectively. Scale bar = 10μm. Zoom scale bar = 1 μm. **B)** ER membrane distribution analysis of VAP-A signals in cells from (A) and (EV5A) using our Fiji binning plugin (described in 2G). Weighted average VAP-A intensity/pixel was compared between bin 1 and bin 3 to acquire a ratio of PN/PP signal distribution. Error bar indicates median and 95% confidence interval (Mann-Whitney U, * p<0,05; ** p<0,01; *** p<0,001; **** p<0.0001). **C)** ER dynamics associated with Vimentin distribution. Representative U2OS cell expressing GFP-Vimentin and ER targeted mCherry-KDEL, imaged by live microscopy. Shown are single channel and overlay frames at t=0 and time color coded (TCC) images from t=0 ranging to t=60, where each of the 60 stacked frames was given a different color LUT. Overlay of all frames shows white areas which were occupied by the ER in every frame, whereas a colorful palette depicts movement of the ER during recording of the video. Cell and nuclear boundaries are demarcated using dashed and continuous lines, respectively. Scale bar = 10μm. Zoom scale bar = 1 μm. **D)** ER dynamics in Vimentin positive or negative regions. Shown are single frame stills from movie S2, showing Vimentin positive (Vim (+)) or (Vim (−)) negative ER every 10 seconds in 5×5μm ROIs. Scale bar = 1 μm. **E)** ER dynamics quantifications of PN versus PP regions. Movement of ER positive pixels in each successive frame in (Vim (+)) or (Vim (−)) of 5×5μm was binned into static (0-60nm/s) or dynamic (61-780nm/s) vectors using membrane displacement analysis (MDA) for Fiji. Error bars indicate median and 95% confidence interval. Quantification was performed on ROIs from 4 cells (Mann-Whitney U test, (**** p<0,0001). **F)** Perinuclear ER dynamics as a function of RNF26 or Vimentin. Stills from live cell movies of representative parental U2OS, Vim KO#2, and siRNF26 cells expressing ER targeted mCherry-KDEL. Shown are single channel images at t=0 and time color coded (TCC) images, where each of 60 frames were color coded and overlaid. Overlay of all frames shows white areas which were occupied by the ER in every frame, whereas a colorful palette depicts movement of the ER during recording of the video. Cell and nuclear boundaries are demarcated using dashed and continuous lines, respectively. Scale bar = 10μm. Zoom scale bar = 1 μm. **G)** Perinuclear ER dynamics quantifications from cells in (F). Movement of ER positive fixels in 5×5μm ROIs was binned into static (0-60nm/s) or dynamic (61-780nm/s) vectors using custom membrane displacement analysis (MDA) for Fiji. Calculations were performed on multiple ROIs from at least 10 cells per condition. Error bar indicates median and 95% confidence interval (Mann-Whitney U, * p<0,05; ** p<0,01; **** p<0.0001).

Since RNF26 and Vimentin form a molecular bridge between the ER and the IF cytoskeleton (Fig. 1), we reasoned that perinuclear ER membranes could be immobilized onto the Vimentin network. Interestingly, overlaying GFP-Vimentin signal onto the ER during time lapse imaging revealed that static ER membranes overlap with the Vimentin cytoskeleton, while Vimentin-free peripheral ER membranes are more mobile (Fig. 4A, movie 2). Furthermore, cells lacking either RNF26 or Vimentin exhibited enhanced mobility of the PN ER (Fig. 4D, EV4C, movies 3-6), indicating that these factors restrict ER network dynamics, and in so doing maintain its spatiotemporal integrity.

### RNF26 and Vimentin mediate PN ER rearrangements during ER stress

The ER adapts its morphology to accommodate a shifting balance of functions during periods of stress. As such, cells treated with tunicamycin to inhibit N-linked glycosylation and subsequent glycoprotein folding, undergo massive expansion in PN ER (Fig. 5A). This so-called ER quality control center (ERQC) enhances the ER’s ability to accommodate accumulating misfolded proteins with the help of ER chaperones, such as Calnexin (Leitman et al., 2013, Leitman, Shenkman et al., 2014). Interestingly, proteolytic lysosomes become recruited to this stress induced PN ER, and the two organelles dwell in association with one another over time (Fig. 5A, B). Reflecting these physiological conditions, PN ER membranes also congregate with Vimentin filaments after tunicamycin treatment (Fig. 5C). Collectively, these observations point towards integration of multiple organellar functions in the PN area during ER stress and suggest a role for Vimentin and RNF26 in this process. Strikingly, ERQC formation appears strongly dependent on the presence of RNF26 and Vimentin, since cells lacking either protein do not accumulate Calnexin in their PN region, instead sustaining a steady state distribution following tunicamycin treatment (Fig. 5D, E and EV5A, B). Importantly, cells also rely on RNF26 and Vimentin for other stress-induced morphological changes of their ER, as inhibition of SERCA-mediated calcium influx by CPA treatment also fails to alter ER morphology under conditions of Vimentin ablation or RNF26 depletion (Fig, EV5C). These results imply that the pairing of Vimentin and RNF26 is necessary to achieve acute morphological changes during ER stress and emphasize the role of the Vimentin IF cytoskeleton in this context.

**Figure 5:**
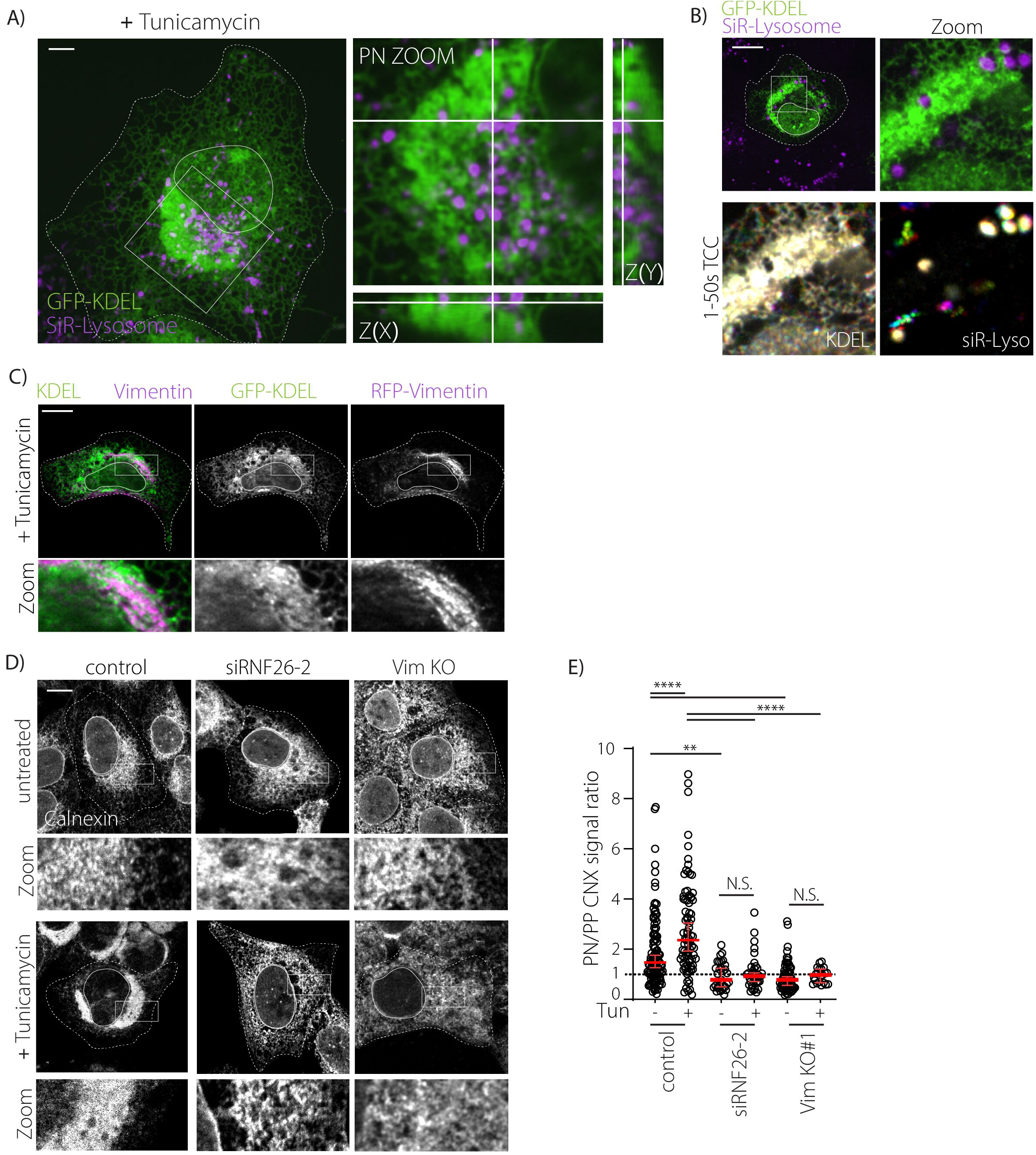
RNF26 and Vimentin regulate compartmentalization of misfolded proteins during ER stress. **A)** Live cell recording of a U2OS cell expressing mCherry-KDEL, treated with tunicamycin (10 μg/mL) to induce ER stress. Lysosomes are visualized with siR-lysosome. Shown is a Z-projection, orthogonal (XZ and XY) views and a tilted zoom box of the perinuclear region. **B)** Live cell recording of a cell as in (A) but followed in time. Shown is a single image at t=0, a zoom in of the perinuclear region and time color coded images of 50 frames of the ER as well as lysosomes. **C)** Live cell image of a tunicamycin-treated U2OS cell expressing RFP-Vimentin and mCherry-KDEL. Shown is an overlay image and single channel images, accompanied with perinuclear zoom boxes. **D)** Confocal analysis of WT U2OS cells, U2OS VIM KO #1 cells, or U2OS cells silenced for RNF26 (si#1), either treated with tunicamycin (5ug/mL) or not, fixed and stained for Calnexin. Shown are representative Z-slices and zoom boxes that show a select region spanning from the nucleus to the PM. **E)** Distribution analysis of Calnexin signal as a function of RNF26, Vimentin and ER stress from images in (A) using our Fiji binning plugin (described in 2G). Weighted average CNX intensity/pixel was compared between bin 1 and bin 3 to acquire a ratio of PN/PP signal distribution. All images show the same magnification. Scale bar = 10μm. All statistical significance tested with Mann Whitney U test. ** p<0,01; *** p<0,001; **** p<0.0001.

## Discussion

Transport and anchorage of organelles in cellular space relies on a multitude of factors, notably including the cytoskeleton and its associated molecular motors. While the roles of microtubule and actin networks with respect to organellar organization and motility have been extensively studied (Reck-Peterson, Redwine et al., 2018, Svitkina, 2018, Zheng et al., 2022), contributions of Vimentin IF in this context are only beginning to emerge (Etienne-Manneville, 2018). In addition to cytoskeletal determinants, physical contacts formed between distinct membrane-enclosed compartments are emerging as important regulators of organellar localization and behavior (Prinz et al., 2020, Wu et al., 2018). How these different modes of spatiotemporal control are integrated to deliver and retain organelles at their site of function remains poorly defined. Here, we report that the ER-embedded RNF26 partners with perinuclear Vimentin intermediate filaments to maintain spatiotemporal integrity of the ER and the endolysosomal cloud. We show that inactive RNF26 employs its cytoplasmically exposed C-terminal tail to bind Vimentin filaments, which accumulate near the nucleus and thus retain RNF26 within the perinuclear ER subdomain. Without this interaction, RNF26 could diffuse throughout the ER and the endosomes would disperse accordingly. Upon activation of the RING domain, RNF26 is released from Vimentin to direct perinuclear positioning of endocytic and proteolytic compartments via ER-endosome contact sites. This interplay between RNF26 and Vimentin IFs helps maintain the bilateral architecture of the ER network and endolysosomal system and supports co-assembly of the ERQC under conditions of ER stress. Our observations thus place a third cytoskeletal component—the intermediate filament—in control of endosomal compartmentalization and the perinuclear ER with implications for ER stress responses.

The Vimentin cytoskeleton has been proposed to serve as a framework for cellular stiffness and provide mechanical support during global rearrangements, such as cell migration (Lowery et al., 2015, Patteson, Vahabikashi et al., 2019, Toivola et al., 2005). In addition, multiple examples of connections between Vimentin and organelles have been reported, including the nucleus (Ketema, Kreft et al., 2013), mitochondria (Tang, Lung et al., 2008), and melanosomes(Chang, Barlan et al., 2009). We now describe a role for Vimentin in maintaining the architecture of the ER and the endolysosomal system. Dispersion of ER membranes from the perinuclear area, as observed here following genetic ablation of Vimentin, has also been reported in response to deregulated microtubules and actin fibers (Gan, Ding et al., 2016, Konietzny, Grendel et al., Noda, Kimura et al., 2014). Furthermore, previous studies have identified ER organizers such as Kinectin, CLIMP63, p180, VIMP, and the long form of STX5, all of which are associated with the microtubule network (Miyazaki, Wakana et al., 2012, Noda et al., 2014, Ogawa-Goto, Tanaka et al., 2007, Shen, Zheng et al., 2019, Vedrenne, 2005). Previously described interplay between Vimentin and microtubule filaments (Gan et al., 2016, Hookway, Ding et al., 2015, Schaedel, Lorenz et al., 2021) or the actin cytoskeleton (Jiu, Lehtimaki et al., 2015, Serres, Samwer et al., 2020) underscores the complexity inherent in the cell’s cytoskeletal foundation. At the same time, our EM reconstruction of the perinuclear region suggests that different types of connections between the ER and various cytoskeletal elements may occur in parallel while we did not observe intense interaction between IF and microtubules. Multiple cytoskeletal networks would allow the ER to harness various ways of fine tuning its architecture and dynamics—and consequently its interactions with partner organelles. Reliance on any one mechanism for perinuclear retention of organelles may however be cell type dependent, as for instance epithelial cells do not appreciably express Vimentin. In these cells the perinuclear region’s integrity may either be simply less important for their physiology or compensated by a stricter control of the microtubule-motor system. In the latter case, continuous active transport through dynein motor activity could be envisioned to ensure perinuclear clustering of organelles (Reck-Peterson et al., 2018).

Considering a rather modest effect of RNF26 depletion on perinuclear ER movement in comparison to that of Vimentin ablation, as well as the vast number of known Vimentin interactors, it is likely that additional ER-associated proteins touch base with the Vimentin cytoskeleton. For instance, Vimentin has been shown to regulate calcium channel distribution in the ER (Dingli, Parys et al., 2012), and IP3Rs have been reported to interact with RNF26 (Fenech et al., 2020), suggesting that IP3R-containing ER membranes may localize to Vimentin or MCS with vesicular calcium stores (Atakpa, Thillaiappan et al., 2018) *through* these joint activities of RNF26. Similarly, the interplay between Vimentin and RNF26 may support transport of LDL-derived cholesterol from late endosomes to the ER for esterification (Sarria, Panini et al., 1992) and perhaps other processes involving endosomal interactions with the ER. From another perspective, reported sensitivity of Vimentin filaments to electrophilic and oxidative reactions (Viedma-Poyatos, Pajares et al., 2020) further opens up the gates for regulation of ER and endolysosomal architecture in response to physiological changes.

Our findings suggest that anchoring of the perinuclear ER on Vimentin intermediate filaments via RNF26 helps maintain the architectural and dynamic integrity of this expansive organelle. Interestingly, we find that Vimentin preferentially binds catalytically inactive RNF26 as compared to its ubiquitination-competent wild type counterpart. This in turn frees active RNF26 to function in endosome positioning through ubiquitin-dependent formation of ER-endosome MCSs (Cremer et al., 2020a, Jongsma et al., 2016). However, restricted localization of RNF26 within the perinuclear ER segment, afforded by its interactions with Vimentin, is also needed to retain the endolysosomal cloud in the corresponding perinuclear cytoplasm. These considerations lead us to propose that RNF26 switches between functional states to sustain dynamic regulation of ER-endosome and ER-cytoskeleton interactions. Within the ER membrane, toggling of RNF26 E3 ligase activity could be accommodated for instance by the availability of its cognate ubiquitin-loaded E2 enzyme UBE2J1 (Cremer et al., 2020a), which is extensively implicated in ER-associated protein degradation (ERAD) (Christianson & Carvalho, 2022, Oikonomou & Hendershot, 2020). Perinuclear integration of lysosomes with ER compartments that accommodate misfolded ER proteins suggest a functional connection between these two organelles. Furthermore, this compartmentalization of misfolded proteins within the ER during ER stress may reflect Vimentin-mediated ‘caging’ previously alluded to the context of cytoplasmic protein aggregates (Johnston, Ward et al., 1998). ER-associated proteotoxic stress can either be cleared by cytosolic proteasomes one molecule at a time in a manner requiring the ERAD machinery (Oikonomou & Hendershot, 2020) or by lysosomes that can receive entire segments of the ER membrane carrying misfolded or aggregated proteins (Fregno & Molinari, 2019). Additionally, excess ER membranes may be removed through direct engulfment by late endosomes upon stress resolution (Loi, Raimondi et al., 2019). In this context, coalescence of endolysosomes at the perinuclear ERQC as afforded by RNF26 and Vimentin may allow for a more efficient stress response.

## Materials and Methods

### Antibodies and reagents

(*Confocal microscopy)* mouse anti-EEA1 (1:200, mAb 610457, BD transduction laboratories), mouse anti-CD63 NKI-C3 (1:500, (Vennegoor & Rümke, 1986), rabbit anti-LAMP1 (1:200, Sino Biological), rabbit anti-VAP-A (1:100, 15272-1-AP, Proteintech), goat anti-CNX, mouse anti-CLIMP63 (followed by anti-Rabbit/Mouse/goat Alexa-dye coupled antibodies (1:400, Invitrogen). SiR-lysosome was used to visualize lysosomes in live microscopy (1:2000, Spirochrome). (*Western blotting)* mouse anti-Vimentin V9 (1:200, SC6260), anti-RFP (Rocha et al., 2009), mouse anti-Vinculin (1:5000, Merck 9131) and rabbit anti HA (1:1000, Cell signaling #3724) followed by secondary Rabbit anti-mouse-HRP, sheep anti-rabbit-HRP (Invitrogen), or goat anti-rabbit or goat anti-mouse IRdye 680 (1:20.000) and IRdye 800 (1:10.000) antibodies (LiCor).

Tunicamycin was bought from Sigma and CPA from Tocris bioscience.

### siRNA transfections

Sequences of the siRNA oligos targeting RNF26 were obtained from Dharmacon (siRNF26-1: CAGGAGGGAUAACCGGAUUUU; siRNF26-2: GAGAGGAUGUCAUGCGGCU). Gene silencing was performed in a 48 or 24 well plate (IF) or 12 well plate (WB) - reagent volumes were scaled up accordingly. In a 24 well plate, 65μL siRNA [50nM] for sequences, see (Jongsma et al., 2016)) was mixed with 26uL 1x siRNA buffer (GE Healthcare) containing 1.15uL Dharmafect 1 transfection reagent. The mix was incubated on a shaker at RT for 40 minutes before the addition of 18.000 HeLa or 30.000 U2OS cells (and coverslips). Cells were cultured for three days before analysis. Non-targeting siRNA or reagent-only was used as a negative control.

### Constructs

RNF26 (and mutants), and GFP-KDEL sequence were expressed from C1/N1 vector series (Clontech) and have been previously described (Jongsma et al., 2016). Vimentin was subcloned between EcoRI and BamHI sites of the C1-IRES-GFP vector of C1-GFP/RFP vector (Clontech). Mutants were created by standard (mutagenesis) PCR methods and sequence verified.

### DNA transfections

Cells were seeded in culture plates to reach approx. 70% confluency on the day of transfection. For IF, cells were transfected with Effectene (Qiagen) (200ng DNA per well of a 24 well plate), according to the manufacturer’s protocol. Larger dishes of HeLa or HEK cells were transfected with PEI at a ratio of 3μg PEI per μg DNA in 200μL DMEM medium. After 15-30 min, the mix was added dropwise to the cells, which were then cultured overnight prior to analysis.

### CRISPR/Cas9-mediated knockout

gRNA sequences targeting the Vimentin gene (see resource table) were cloned into the BbsI site of PX440 (containing the Cas9 gene and a puromycin resistance gene). This plasmid was transfected into HeLa or U2OS cells and the next day, cells were selected with 200ug/mL puromycin for 3 days. Then, cells were diluted and cultured in a 15cm dish, allowing separated colonies to grow. These were isolated, expanded, and analyzed for loss of Vimentin by Western blot.

### Immunofluorescence confocal microscopy

Cells grown on coverslips (Menzel Gläser) were fixed with 3.7% paraformaldehyde, washed three times with PBS, permeabilized with 0.1%TX100 for 10 min and blocked in 0.5% BSA for one hour. Next, coverslips were incubated with primary antibodies in 0.5% BSA for 1hr at RT, washed and incubated with Alexa-labeled anti mouse/rabbit/rat secondary fluorescent antibody or streptavidin. After washing, coverslips were mounted on glass slides with ProLong Gold with DAPI (Life Technologies). Samples were imaged with a Leica SP8 confocal microscope equipped with appropriate solid-state lasers, HCX PL 63 times magnification oil immersion objectives and HyD detectors (Leica Microsystems, Wetzlar, Germany). Data was collected using 2048 x 2048 scanning format with line averaging without digital zoom, or 1024 x 1024 scanning format with digital zoom in the range of 1.0-2.0 with line averaging. Images were smoothened with 1x pixel average filter in ImageJ. Overlap analysis was performed using JaCoP analysis tool in Fiji with manual threshold selection. Near-superresolution images were acquired with a Zeiss Airyscan microscope with 100x objective, 2x pixel sampling and deconvolved using Zen software.

### Endosomal distribution analysis

To analyze endosomal distribution, we developed a Fiji plugin that segments the cytoplasm into areas of varying distance to the nucleus. In short, the plugin uses thresholding to segment the nuclei, followed by optional manual correction. The outer edges of the cell are manually annotated. Then, using a Euclidian distance map, we measure for each pixel within the cytoplasm (outside nucleus and within the cell) the relative (normalized to the maximum distance per cell) distance to the nucleus. These distance values are then segmented into a selectable number of bins, resulting in the segmentation shown in Fig. 2G.

### Live microscopy

For live cell microscopy, cells were seeded in 4-chamber live cell dishes and imaged under conditions of 37°C and 5% CO_2_ with a Leica SP8 confocal microscope equipped with solid state lasers or a Andor Dragonfly spinning disc module. Data was collected using 63x or 100x oil immersion objectives in a 2048×2048 scanning format at 3 frames/sec. Time color coded images were made with Fiji.

### ER membrane dynamics analysis

ER membrane displacement was quantified using a custom developed Fiji plugin that has been described earlier (Spits et al., 2021). The method is based on optical flow analysis, which defines corresponding areas between frames and generates a vector map. Live cell movies were simplified by removing every second and third frame, leaving a 1 frame/sec video for analysis. Membrane displacement (60nm/sec) was binned into 7 displacement bins, ranging from 0 to 60 nm/s (resulting in displacement bins of nm/s). The first two bins (i.e., 0 and 60nm/s displacement) were classified as ‘static ER’. ROIs of 5×5um from at least 10 cells were compared across conditions, unless indicated otherwise.

### Proximity biotinylation pulldowns

HeLa cells transfected to express proteins of interest and were treated as indicated in the figure legends before three washes with PBS and subsequent cell lysis in 0.5% TX-100 lysis buffer (150mM NaCl, 50mM Tris-HCl pH 7.5, 5mM MgCl2, 0.5% (v/v) TX-100 and protease inhibitors (Roche diagnostics, EDTA free)). Nuclei were lysed by adding 1:4 SDS buffer (2% SDS, EDTA) and samples were sonicated (5×1s pulses, 80% power, Fisher Scientific). Samples were diluted to 0.2% SDS with TX-100 lysis buffer and centrifuged for 20 min at 20,000 x g. After spinning, samples were incubated with Neutravidin-agarose beads (Pierce) overnight at 4oC. Beads were washed 4 times with 1% SDS in PBS before elution in a 2x Laemmle sample buffer by boiling for 10 min and SDS-PAGE.

### SDS-PAGE and Western blotting

Samples were separated by 10% SDS-PAGE and transferred to nitrocellulose in ethanol-containing transfer buffer at 300mA for 2-3 h. The membranes were blocked with 5% milk/PBS before incubation with primary antibody diluted in blocking buffer for 1hr at RT. After washing twice with PBS/0.1% Tween-20, proteins were detected with secondary antibodies. Fluorescent signals were directly imaged with an Odyssey Fx laser scanning fluorescence imager (Li-COR Biosciences) and quantified using LiCor ImageStudio software.

### Electron microscopy and tomography

#### 3D tomography and rendering

Adherent U2OS cells cultured in a 5 cm diameter Petri dish were fixed for 2 hours at room temperature in 0.1M Cacodylate buffer containing 1,5% glutaraldehyde. The fixed cells were incubated for 1 hour on ice in a Cacodylate buffer containing 1% Osmium tetroxide and subsequently in water containing 1% uranyl acetate. The cells were then dehydrated through a series of ethanol steps and embedded in Epon. The flat embedded cells were sectioned with an ultramicrotome using a diamond knife at a nominal section thickness of 200 nm. The sections were transferred to a formvar and carbon coated 0.5×2 mm copper slot grid and stained for 20 minutes with 7% uranyl acetate in water and for 10 minutes with lead citrate according to Reynolds (Reynolds, 1963). Electron microscopy images of serial sections were recorded by a Tecnai 20 TEM (Thermo Fisher Scientific) equipped with an EAGLE 4×4K digital camera. The tilt series for electron tomography were collected using Xplore3D (Thermo Fisher Scientific) software. The angular tilt range was set from −60° to 60° with 2° increments. Alignments of the tilt series and weighted-back projection reconstructions for tomography were performed using IMOD (Kremer, Mastronarde et al., 1996). The drawing and annotation of subcellular structures in the tomograms and the alignment of the serial tomograms was done manually using AMIRA (Thermo Fisher Scientific). Generated surfaces were exported to Cinema 4D (Maxon), which was used to smooth the surfaces, and to render images and movies (Redshift, Maxon).

#### Immuno-EM

For immunogold-EM, U2OS cells transfected as indicated were prepared for cryosectioning, as described (Van Elsland et al., 2016; Peters et al., 2006). Briefly, cells were fixed for 24 h in freshly prepared 2% paraformaldehyde in 0.1M phosphate buffer. Fixed cells were scraped, embedded in 12% gelatin (type A, bloom 300, Sigma) and cut with a razor blade into 0.5 mm3 cubes. The sample blocks were infiltrated in a phosphate buffer containing 2.3M sucrose. Sucrose-infiltrated sample blocks were mounted on aluminum pins and plunged in liquid nitrogen. The vitrified samples were stored under liquid nitrogen. Ultrathin cell sections of 75nm were obtained essentially as described elsewhere (van Elsland, Bos et al., 2016). Briefly, the frozen sample was mounted in a cryo-ultramicrotome (Leica). The sample was trimmed to yield a squared block with a front face of about 400 x 300 μm (Diatome trimming tool). Using a diamond knife (Diatome) and antistatic devise (Leica) a ribbon of 75nm thick sections was produced that was retrieved from the cryo-chamber with the lift-up hinge method (Bos et al., 2011). A droplet of 2.3M sucrose was used for section retrieval. Obtained sections were transferred to a specimen grid previously coated with formvar and carbon. Grids containing thawed cryo-sections of fixed cells were incubated on the surface of 2% gelatin at 37°C. Sections were then rinsed to remove the gelatin and sucrose, were blocked with 1% BSA in PBS, labeled with the indicated antibodies and 10/15 nm protein A-coated gold particles (CMC, Utrecht University) and embedded in 1.8% methylcellulose and 0.6% uranyl acetate. Electron microscopy imaging was performed with a Tecnai 12 transmission electron microscope (FEI) operated at 120 kV acceleration voltage.

### Statistics

All error bars in WB quantification correspond to SD or the mean from three independent experiments. For microscopy, cells from at least two independent experiments were used. In distance bin analysis, error bars correspond to the median of measurements with 95% confidence interval. Statistical evaluations report on Student’s t-test (for gel and colocalization analysis) with error bars corresponding to mean and standard deviation, or Mann-Whitney U non-parametrical test (for membrane distribution and displacement analysis), with error bars corresponding to median and 95% confidence interval, as described in the legends,. *p,0.05, **p<0.01, ***p<0.001, ****p<0,0001; ns: not significant.

## Acknowledgements

We thank M.L. Jongsma, L. Janssen, and J.J. Akkermans for DNA constructs and valuable discussions. This work was supported by an ERC Adv grant (ERCOPE) to J. Neefjes.

## Author contributions

TC, IB, and JN designed the study. TC performed most of the experiments, supported by LtH. Immuno-EM was performed by DvE, tomography and EM by EB and reconstruction of the images by RK. Fluorescence image analyses and software programming was performed by LV. TC, IB, and JN wrote the manuscript with input from all authors. JN and IB coordinated the study.

## Supporting Information

Expanded view Figures PDF

Movies EV1-6

**Fig. EV1: RNF26 is immobilized in the ER membrane its inactive state**

**A)** Intracellular distribution of organelles. EM micrograph of U2OS cell with boxed regions indicating perinuclear (PN) and peripheral (PP) zoom-ins. Scale bar = 10 μM.

**B)** Zero tilt EM image of zoom-in region designated in (A), showing region used for tomography in (1C). Pseudo colored annotation of the ER (cyan), mitochondria (red), lysosome (green) Golgi apparatus (magenta) and (in zoom box) intermediate filaments (yellow) as manually drawn in. Scale bar = 1μM.

**C)** Examples of IF-ER connections from select regions in (B) and adjoined serial sections used for tomography. Arrows indicate Vimentin IFs (yellow) and ER structures (cyan). Scale bar = 100nm.

**D)** Contribution of pseudocompatible UBE2J2 to the interaction between RNF26 and Vimentin. Proximity biotinylation of RNF26 I382R or RNF26 WT in HEK293 alone or in coexpression of RFP-UBE2J2 or EV plasmids. Cells were treated with 250μM biotin prior to lysis, Membranes were stained for endogenous Vimentin, HA and RFP. Molecular weight markers indicated (MW). Representative scan of three biological replicates.

**E)** RNF26 mobility in ER membrane as a function of RING presence or activity. Fluorescence recovery after photobleaching (FRAP) of RFP signal in HeLa cells expressing RFP-RNF26 WT, I382R or ΔRING. Signal was bleached to approximately 20% of original before imaging of signal recovery every 3 sec for 50 frames.

**F)** Quantifications of FRAP in (E), showing average signal recovery of RNF26 WT, I382R or ΔRING. Scatter plot shows mean +/− SD of total recovery at endpoint (t=150s) in 13 (WT), 8 (I382R) and 8 (ΔRING) cells (T-test. ** p<0,01; *** p<0,001; **** p<0.0001).

**Fig. EV2. Vimentin is essential for filamentous appearance of RNF26 I382R**

**A)** RNF26 I382R distribution as a function of Vimentin. Confocal microscopy images of U2OS and U2OS VIM KO#1 cells transiently transfected to express RFP-RNF26 I382R were fixed and immunostained for VAP-A. Shown are single channel RFP images, and colored overlay images. Boxed zoom-ins show perinuclear magnification.

**B)** Cell and nuclear boundaries are demarcated using dashed and continuous lines, respectively. Scale bar = 10μm.

**C)** Validation of Vimentin knockout in HeLa cells. Western blot analysis of WT and Vimentin knockout Hela cells created using CRISPR/Cas9. Membranes were stained for Vimentin, while Tubulin was used as a loading control. Molecular weight markers indicated.

**D)** Rescue of RNF26 I382R distribution in Vimentin knockout cells. Confocal microscopy images of HeLa VIM KO cells transfected with RFP-RNF26 I382R and Vimentin-IRES-GFP before fixing and imaging. Shown is an overlay image of GFP and RFP-RNF26 IR signal and zoom-ins of a 1) GFP negative (VIM KO) cell and 2) GFP positive (rescued) cell. Scale bar =

**Fig. EV3: ER Localization and activity of RNF26 mutants**

**A)** Confocal analysis of mutants described in (3A). U2OS cells were transfected with indicated constructs, fixed for IF and labeled for VAP-A before imaging with a confocal microscope. Images show a single Z-slice of a representative cell. Cell and nuclear boundaries are demarcated using dashed and continuous lines, respectively. Scale bar = 10μm.

**B)** Ubiquitination of RNF26 WT versus RNF26 I382R or RNF26 Y432A. HeLa cells were transfected with RFP-tagged RNF26 constructs and HA-Ubiquitin before RFP immunoprecipitation under denaturing conditions. Samples were separated by SDS-PAGE and Western blotting. Membrane was stained for RFP and Ub. Shown are single channel images and overlay images of RFP (green) and Ub (red). Position molecular weight standards indicated.

**Fig. EV4: RNF26 and Vimentin regulate VAP-A, CLIMP63 and Calnexin distribution**

**A)** Overflow confocal images of experiment in (4A), showing VAP-A signal in another VIM KO cell line and U2OS cells treated with a second siRNA oligo. Cell and nuclear boundaries are demarcated using dashed and continuous lines, respectively.

**B)** ER sheet distribution as a function of RNF26 or Vimentin. Confocal analysis of parental U2OS cells, U2OS VIM KO #2 cells, or U2OS cells silenced for RNF26 (si#2) that were fixed and immunostained for CLIMP63 prior to imaging with a confocal microscope. Shown are Z-slices of representative cells. Zoom boxes show a select region spanning from the nucleus to the PM. Cell and nuclear boundaries are demarcated using dashed and continuous lines, respectively. Scale bar = 10μm. Zoom scale bar = 1 μm.

**C)** Overflow live cell images of perinuclear ER dynamics in siC cells (4G). Cell and nuclear boundaries are demarcated using dashed and continuous lines, respectively. Scale bar = 10μm. Zoom scale bar = 1 μm.

**Figure EV5: RNF26 and Vimentin regulate ER morphology during ER stress**

**A)** Confocal analysis of WT HeLa cells or WT HeLa, HeLa VIM KO #1 cells or HeLa cells silenced for RNF26 (si#1) that were treated with tunicamycin or that were fixed and stained for calnexin. Shown are Z-slices of representative cells and zoom boxes that show a select region spanning from the nucleus to the PM.

**B)** Distribution analysis of Calnexin signal as a function of RNF26, Vimentin and ER stress from images in (A) using our Fiji binning plugin (described in 2G). Weighted average CNX intensity/pixel was compared between bin 1 and bin 3 to acquire a ratio of PN/PP signal distribution. Statistical significance was tested using Mann-Whitney U test. All statistical significance tested with Mann Whitney U test. ** p<0,01; *** p<0,001; **** p<0.0001.

**C)** Confocal analysis of WT U2OS cells, U2OS VIM KO #1 cells, or U2OS cells silenced for RNF26 (si#1), either treated with cyclopiazonic acid (CPA) (25μM) or not, fixed and stained for Calnexin or VAP-A. Shown are representative Z-slices.

All images show the same magnification. Scale bar = 10μm.

**Suppl. Movie 1**: Fly-over movie of 3D TEM tomogram of a perinuclear region of a U2OS cell, accompanying Fig. 1. Zoom ins are accompanied by original TEM micrographs, showing close connections of the ER and intermediate filaments. Ultrastructures were manually annotated and pseudocolored as following: ER (cyan), mitochondria (red), lysosome (green) Golgi apparatus (magenta), intermediate filaments (yellow) and microtubuli (grey).

**Suppl. Movie 2**: Live cell recording of a U2OS cell transiently expressing GFP-Vimentin (green) and mCherry-KDEL (magenta). Shown is an overlay of both channels over 60 frames (1 frame/sec). Stills, zooms, and time color coded image are shown in Fig. 4F.

**Suppl. Movies 3-6**: Live cell recording of ER dynamics as a function of Vimentin or RNF26. Shown are 60 sec (3 frames/ sec) movies of U2OS WT (Movie 3), VIM KO#1 (Movie 4), siC-treated (Movie 5) or siRNF26 (si#1)-treated (Movie 6) cells expressing mCherry-KDEL.

